# AML patient blasts exhibit polarization defect upon interaction with bone marrow stromal cells

**DOI:** 10.1101/2024.06.21.600083

**Authors:** Khansa Saadallah, Benoît Vianay, Louise Bonnemay, Hélène Pasquer, Lois Kelly, Cécile Culeux, Raphael Marie, Sofiane Fodil, Paul Chaintreuil, Emeline Kerreneur, Arnaud Jacquel, Emmanuel Raffoux, Rémy Nizard, Laurent Blanchoin, Lina Benajiba, Manuel Théry

## Abstract

Hematopoietic stem and progenitor cells (HSPCs) establish specific interactions with bone marrow stromal cells, leading to their polarization. Given the role of cell polarity in protection against tumorigenesis and the importance of the niche in hematological disorders such as acute myeloid leukemias (AMLs), we investigated the polarization capacities of leukemic blasts from patients. Using engineered micro-niches and centrosome position with respect to the contact site with stromal cells as a proxy for cell polarization, we showed that AML cell lines and primary cells from AML patient blasts were unable to polarize in contact with healthy stromal cells. In return, exposure to AML patient-derived stromal cells compromised the polarization of healthy adult HSPCs and AML blasts from patients. Using live cell imaging in engineered “bone-marrow-on-a-chip”, we further revealed that stromal cells from a leukemic niche increased the migration speed and distance of healthy HSPCs and AML blast as compared to their behavior in contact with healthy stromal cells. The results collectively demonstrated the respective influences of intrinsic AML blast transformation and extrinsic contact with AML stromal cells on the defective polarization of AML blast. They suggested that leukemic progression is associated with cell polarization defects and proposed new methodological approaches to investigate this relationship in AML progression.

## Introduction

Acute myeloid leukemias (AMLs) constitute a diverse group of clonal hematopoietic disorders characterized by the accumulation of abnormal yet highly proliferative myeloid progenitor cells referred to as blasts^1,2^. The sustained self-renewal and extensive proliferation of the blast population are fundamental aspects contributing to the persistence of leukemia^3^. AMLs malignancies ultimately lead to failure of hematopoiesis by disrupting the normal commitment of hematopoietic stem cells (HSCs) and progenitor cells towards myeloid differentiation resulting in cytopenia and debilitating complications^4^ contributing to the mortality associated with AMLs. The genesis of this hematological malignancy stems from a confluence of environmental and genetic modifications. It is widely recognized as a clinically heterogeneous disease with significant variability in post-treatment survival influenced by factors such as age, blast cell morphology, cytogenetic abnormalities and gene mutations^5^.

There is speculation that leukemic cells hijack and disrupt the supportive microenvironment of HSCs through several potential targets exploited by leukemic stem cells (LSCs) including adhesion molecules (CD44) or chemokines receptors (CXCR4)^6^. This phenomenon has the potential to shift the balance of microenvironments from supporting steady-state hematopoiesis to conditions favoring accelerated expansion of leukemic cells, potentially contributing to leukemogenesis^7^ and chemoresistance development. The characterization of both healthy and diseased hematopoietic cells is inextricably intertwined with the dynamic compartment in which they develop, the bone marrow niche. Consisting of stromal (mesenchymal stem cells (MSCs), osteoblasts, endothelial cells (ECs)…) together with hematopoietic cellular components, the hematopoietic niche exerts a substantial influence on hematopoiesis, the development, and/or maintenance of leukemia, not to mention the alteration of the niche itself in the context of leukemic transformation^8^. It is therefore imperative to understand the fundamental role of the microenvironment in the progression of the disease to gain a comprehensive understanding of the intrinsic communication between all components within the hematopoietic niche. Unfortunately, the complex interplay of chemical and physical factors in the niche makes it challenging to discern the specific contributions of stromal and hematopoietic cells to the development of AML.

In various tissues, maintaining the balance between stem cell expansion and differentiation relies on the regulation of cell polarity and asymmetric cell divisions (ACD)^9,10,11,12,13,14,15^. There is growing evidence to support the tumor-suppressive roles of polarity proteins in mammals, in particular their impact on cell-to-cell or cell-to-matrix interactions^16,17,18^. HSCs located at the apex of normal hematopoiesis, undergo polarization during migration, involving shape alterations and protein segregation. It has been elucidated that intercellular contact led to HSC polarization by inducing the formation of a cellular protrusion called magnupodium^19,20,21^. This protrusion formation occurs alongside the polarization of certain proteins, such as CD133 (Prominin-1)^22^ and CD44^23^. Consistent with *in vivo* observations^24^, our laboratory has recently found that HSCs and HSPCs manifest a variety of shapes, with a predominant elongated morphology observed during interactions with osteoblasts or ECs^25^. HSPC appeared capable of polarizing in contact with stromal cells, as revealed by the position of their centrosomes toward the magnupodium they formed in contact with stromal cells^25^. This opens the question of the ability of leukemic blasts to polarize in contact with stromal cells and the role of this polarization in the maintenance of proper hematopoiesis or AML development.

Although the influence of polarity on hematopoietic stem cell function is a relatively new field, it is gaining traction, especially in the context of hematopoietic malignancies such as leukemias. The relevance of polarity in disease initiation, progression, or suppression, is intricately linked to an understanding of leukemia’s origin. For instance, inhibition of Llgl1, a component of the Scribble complex known for its role in HSCs renewal, diminishes survival in AMLs^26^. The inactivation of Scribble partners induces dysregulation in the proliferation, signaling, and motility of HSCs. Also, the disruption of Yap1/Taz co-polarization with Cdc42 through the Scribble complex precipitates the loss of quiescence and self-renewal in HSCs^27^. Furthermore, there is evidence indicating a loss of polarity in HSCs during the aging process, characterized by depolarization of Cdc42^28^. The restoration of polarity through Cdc42 depletion facilitates the differentiation of AML cells^29^. Conversely, polarity alterations can prompt the redistribution of pro-differentiation cues, consequently suppressing tumorigenesis. Therefore, inhibition of Lis-1, a regulator of mitotic spindle positioning, disrupts Numb segregation inducing its polarization, thereby fostering the differentiation of AML cells^30^. These observations underscore the importance of investigating polarity, as it is intricately linked to mechanisms compromised in the leukemic context. The interaction between hematopoietic cells and niche cells assumes a critical role in preserving hematopoietic integrity.

This study employs engineered artificial niches at two different scales to investigate communication between healthy or AML bone marrow stromal cells and HSPCs or blasts from AML cell lines or patients. The first method consists of a confined culture model of cell-sized microwells constraining a pair of a stromal cell with a hematopoietic cell allowing prolonged cell-cell interactions whilst restricting migration. The second method involves a Bone Marrow on Chip (BMoC)^31^, which is proposed as a microfluidic model for three-dimensional coculture promoting both migration and interaction.

## Results

To investigate the functional impact of hematopoietic cells and stromal cell interactions, polyacrylamide microwells coated with fibronectin *(Figure1.A-B)* were employed to isolate these cells for a 24-hour interaction period, followed by fixation and immunofluorescence. Centrosome positioning was utilized as a polarity marker, and the quantitative analysis of its localization regarding the contact region with stromal cells served as a measurement of the cell polarization index (PI) (*Figure 1.C*).

**Figure 1:**
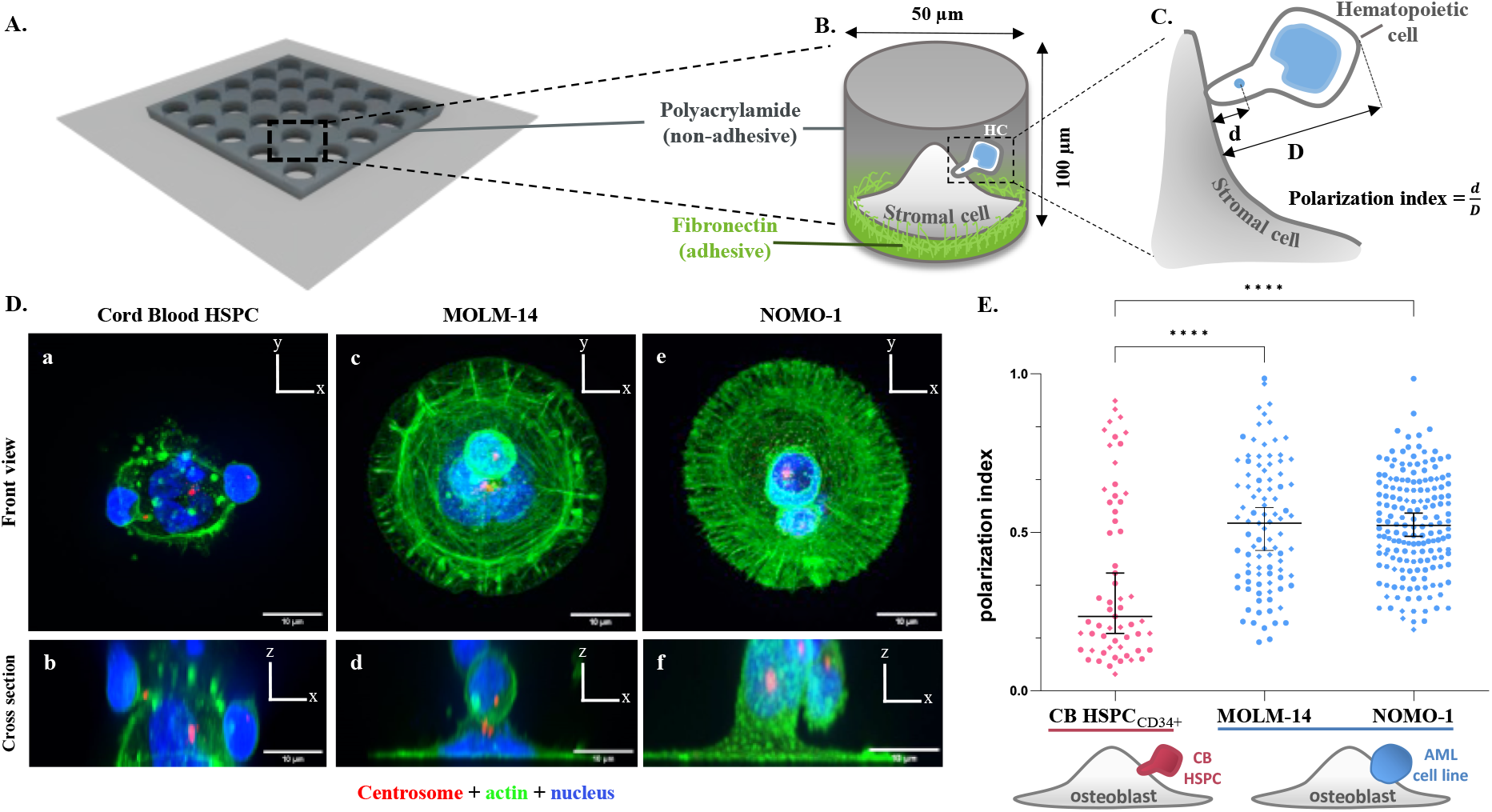
AML cell lines lack polarization when interacting with osteoblasts in comparison to polarized cord blood (CB) HSPCs within microwells. **(A)**.Polyacrylamide microwells on a cover slide. **(B)**. Zooming in on a singular microwell that illustrates the distribution of the non-adhesive (polyacrylamide) and the adhesive part (fibronectin coating) on the bottom, facilitating the spreading of stromal cells. The microwells have diameters of 50 µm and a depth of 100 µm, designed to prevent cell escape during the culture **(C)**. Further magnification focuses on the interactions occurring within the wells, allowing for the measurement of the polarization index (d/D). **(D)**. Polarization defects in AML cell lines interacting with osteoblasts (hFOB) within microwells. **(a**,**b)**. CB HSPCs display magnupodia where can be localized their centrosomes (red). **(c**,**f)**. MOLM-14 and NOMO-1 leukemic blasts are both round and do not exhibit magnupodia upon interaction with the osteoblast. The shape of the cell is visualized via actin labelling shown in green. Cells nuclei are in blue. **(E)**. Quantification of the polarisation index of AML cell lines, MOLM-14 and NOMO-1 in contact with osteoblasts (hFOB) in comparison to CB HSPCs within microwells. Each spot shape denotes a biological replicate (two or three replicates; n _CB HSPCs_ =59, n _MOLM-14_ =95; n _NOMO-1_ =173). The mean of the medians is represented by a black bar. Differences between populations were evaluated using non-parametric test Kruskal-Wallis ANOVA with a P-value <0.0001 (****).

Pairs of osteoblasts (hFOBs) and cord blood-derived (CB) CD34^+^HSPCs were confined in microwells *(Figure 1.D.a-b)*, and their interactions were compared with those involving AML leukemic cells from MOLM-14 and NOMO-1 cell lines *(Figure 1.D.c-f)*. Notably, CB HSPCs demonstrated significant polarization *(Figure1.E)* with a distinctive magnupodia pseudopod formation upon attachment to osteoblasts aligning their centrosome toward the contact zone *(Figure 1.D.a-b)*. Conversely, both MOLM-14 and NOMO-1 blasts predominantly exhibited a symmetrical, rounded morphology *(Figure 1.D.c-f)* with random polarization during their interactions with osteoblasts *(Figure 1.E*). However, cell lines have acquired specific characteristics associated with their adaptation to culture conditions, potentially deviating from the behavior exhibited by blasts within the bone marrow. The prolonged absence of stromal cells in their environment may compromise their anchorage machinery. Alternatively, defective anchorage and polarization mechanisms could represent a distinctive feature of AML progression. To further investigate these possibilities, it was necessary to work with primary cells derived directly from patients.

Primary cells, specifically bone marrow-derived MSCs (MSCs) and bone marrow-derived CD34^+^-HSPCs, were sourced from healthy donors (HD) undergoing hip surgery for prosthesis placement. For the malignant counterpart, primary bone marrow derived MSCs (AML MSCs) and blasts were harvested from AML patients. AML cells were obtained from individuals with distinct mutational statuses (*supp. Table.1*) displaying high percentages of blasts from patients with an age range of approximatively 50 years (aged between 20 and 70), facilitating a meaningful comparison between them while mitigating the influence of age-related biases (supp. *Table.2*). The cell collection was performed at the time of diagnosis, prior to any treatment or chemotherapy. Thus, freshly collected AML patient blasts and healthy HSPCs were promptly plated into the engineered niches alongside stromal cells.

The interaction dynamics between adult HSPCs and osteoblasts (from osteoblastic cell line) (*Figure 2.A.a-b*) or HD MSCs (primary cells) (*Figure 2.C.a-b)* in microwells unveiled their capability of magnupodium formation directed toward the contact zone. Notably, blasts from all three AML patients mainly exhibited rounded shapes when in contact with the healthy stromal cell, devoid of the distinctive presentation of a narrow pseudopod observed in adult HSPCs (*Figure 2.A.c-h, C.c-h*).

**Figure 2:**
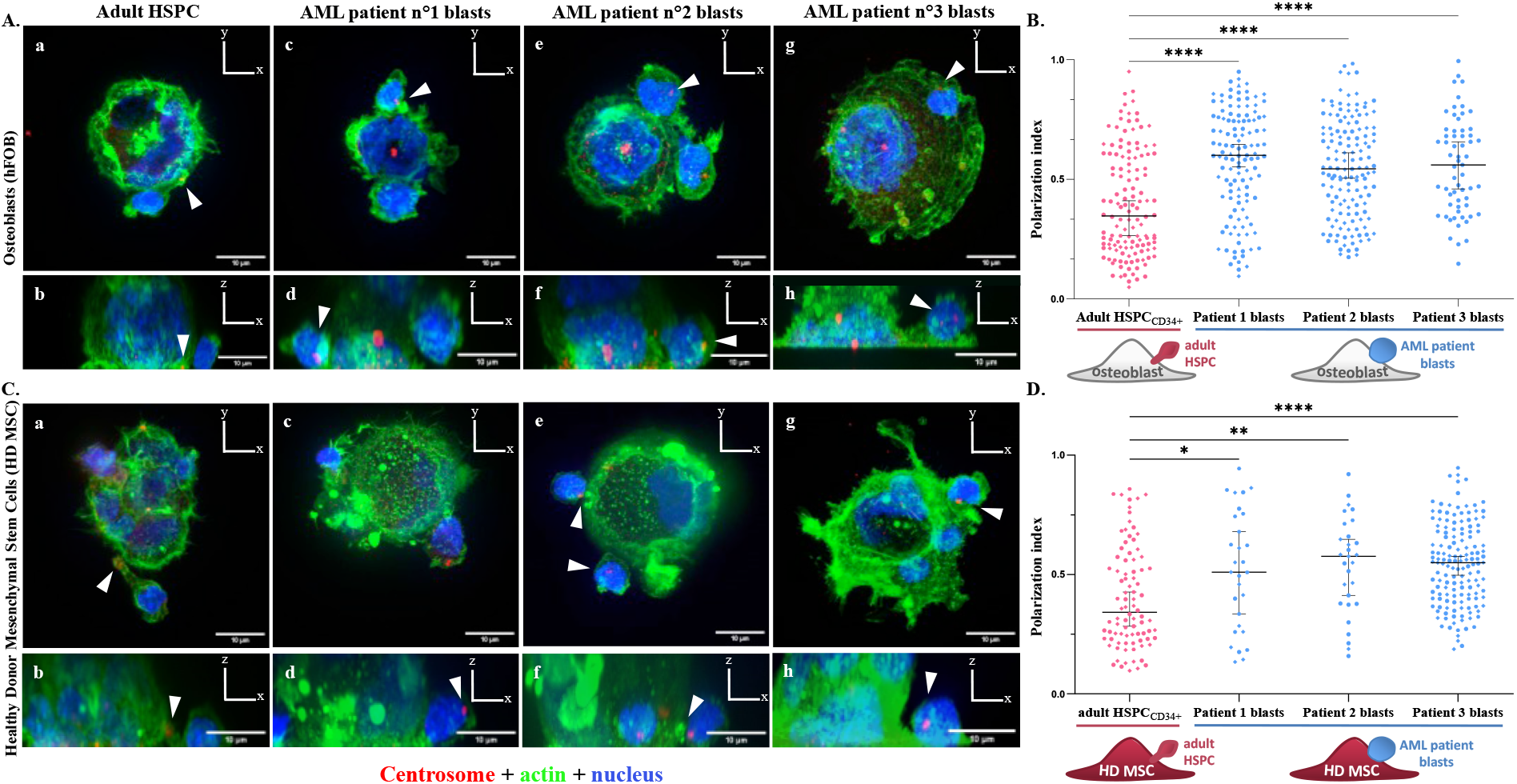
AML patient blasts exhibit impaired polarization when interacting with distinct bone marrow stromal cells. **(A and C)**. Adult HSPCs, AML primary blasts from patients 1, 2 and 3 blasts interacting with osteoblasts (hFOB) (A) or healthy donor MSCs (HD MSCs) (C) in microwells. **(a**,**b)**. Similar to CB HSPCs, adult HSPCs elongate to display magnupodia where can be located their centrosomes (red, indicated by arrows) in close vicinity to the contact site with both stromal cell types. **(c-h)** AML primary blasts from patients 1, 2 and 3 exhibit large zones of contact with no sign of magnupodia upon interaction with the stromal cell (osteoblast or HD MSC). The shape of the cells is visualized via actin labelling shown in green. Cells nuclei are in blue. **(B and D)**. Quantification of the impaired polarisation index of AML patient blasts, compared to adult HSPCs, in contact with osteoblasts (hFOB) (B) or HD MSCs (D) within microwells. **(B).** AML primary blasts from patients 1, 2 and 3 display a random distribution of their centrosome positioning while interacting with osteoblasts. Each spot shape denotes a biological replicate (three replicates; n _adult HSPCs_ =132, n _patient 1 blasts_ =123; n _patient 2 blasts_ =142, n _patient 3 blasts_=61). The mean of the medians is represented by a black bar. Differences between populations were evaluated using a Kruskal-Wallis ANOVA test with P-values <0.0001 (****). **(D)**. AML primary blasts from patients 1, 2 and 3 display a random distribution of their centrosome positioning while interacting with HD MSCs. Each spot shape denotes a biological replicate (three replicates; n _adult HSPCs_ =88, n _patient 1 blasts_ =148; n _patient 2 blasts_ =29, n _patient 3 blasts_ =29). The mean of the medians is represented by a black bar. Differences between populations were evaluated using a Kruskal-Wallis ANOVA test with P-values of 0.0301, 0.0045 and <0.0001 (*, ** and ****).

Furthermore, in a manner analogous to the description of cord blood derived HSPCs (CB-HSPCs), adult CD34^+^-HSPCs exhibited a preserved ability to orient their centrosome towards osteoblast (*Figure 2.B*) and healthy bone marrow stromal cell (*Figure 2.D*). In contrast, the three AML blasts from patient donors displayed a random distribution of their centrosome localization in all conditions. This indicated a lack of polarization of AML patient blasts toward their contact site with osteoblasts (*Figure 2.B*), and with bone marrow HD MSCs (*Figure 2.D*). Thus, the polarization mechanism appeared functional in healthy adult HSPC and intrinsically perturbed in AML blast. However, the polarization process is contingent upon the interplay between diverse cell types, so it might also be regulated by the stromal cells. The AML niche being instrumental to tumoral development, it appeared necessary to specifically test the potential role of stromal cells from AML patients on the polarization of healthy HSPC and leukemic blasts.

To that end, we first tested the ability of HSPCs derived from healthy donors (HD HSPCs) to polarize in contact with MSCs derived from healthy donors or AML patients (Figure *3.A.c-f*). Interestingly, HD HSPCs from the same donor that polarized in contact with HD MSCs (*Figure 2.D*.), displayed a random orientation of their centrosomes in contact with AML MSCs (*Figure 3.B*.). This observation demonstrated that transformed stromal cells could impair the polarization of healthy HSPC, although the machinery underlying the control of centrosome position was functioning properly in these cells. Furthermore, when mimicking the conditions within the bone marrow of patients, ie by seeding patient-derived blasts onto AML-derived stromal cells, we observed that the AML patient blasts not only lacked the ability to polarize in contact with AML MSCs but even exhibited a tendency to polarize in the opposite direction (*Figure 3.A.g-h*.). The majority of centrosomes were positioned toward the uropod, situated at the rear of the cell, as shown by the values of polarity indices between 0.5 and 1 (*Figure 3. B*.). This orientation of polarization toward the rear of the blasts was reminiscent of the polarization observed in migrating leukocytes^32,33^. This suggested that the observed inverted polarization of AML blast on AML-derived stromal cells may result not only from defective anchorage but also from the stimulation of their migration properties.

**Figure 3:**
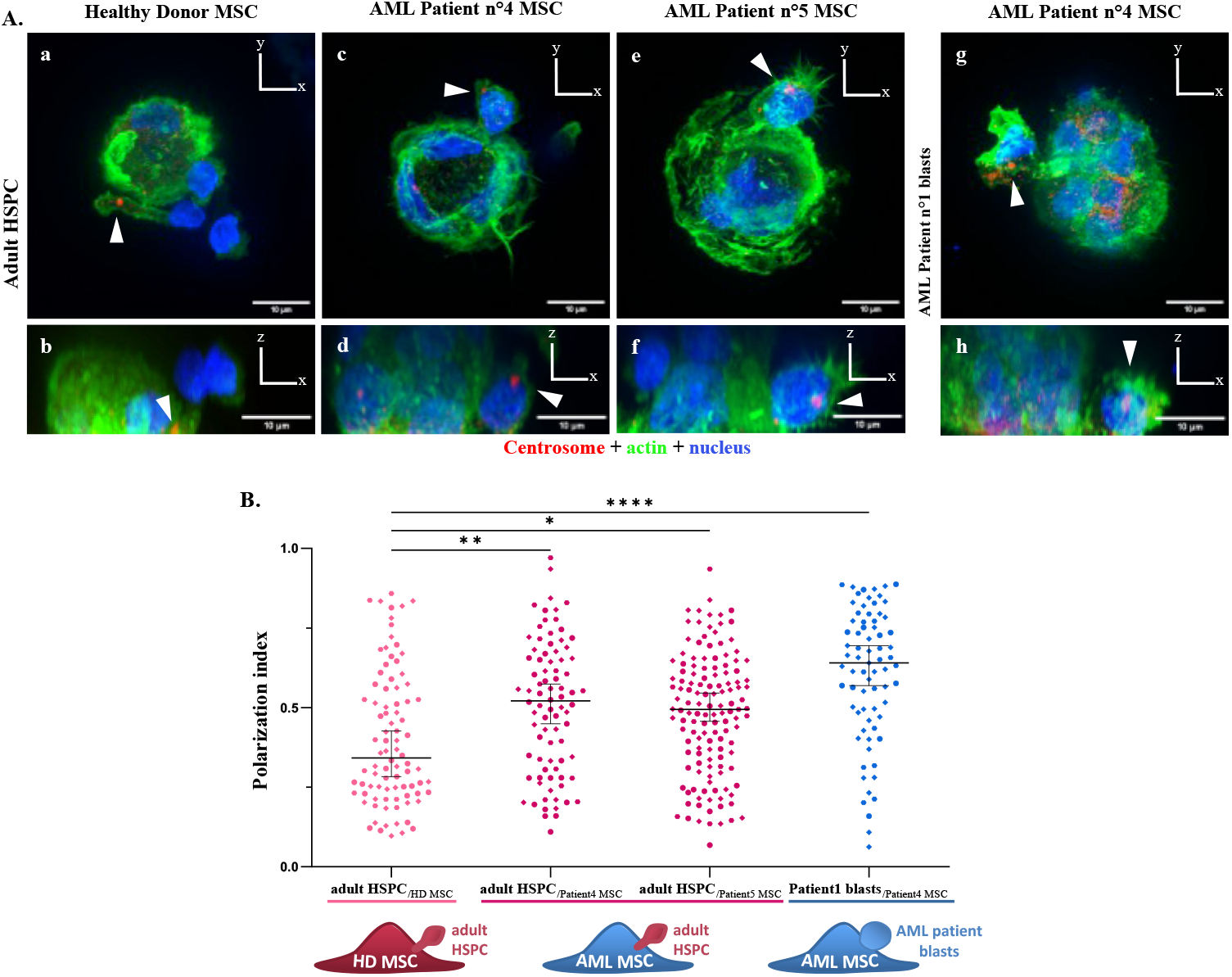
The nature of the niche impacts the adult HSPC’s ability to form magnupodia and elongate upon interaction with either HD or AML MSC in comparison to a full leukemic microwell containing AML primary patient blast interacting with AML MSC. **(A).** Adult HSPCs interacting with either healthy or AML bone marrow stromal cell. **(a**,**b)**. Polarized adult HSPCs interacting with HD MSCs. **(c-d)** Adult HSPCs interacting with AML MSCs derived from patient 5 **(c**,**d)** and 4 **(e**,**f)**. Cells are mainly exhibiting more random centrosome positions **(g-h)**. AML patient blasts interacting with AML MSCs from patient 4. Centrosome is indicated in red and marked by an arrow. The shape of the cells is visualized via actin labelling shown in green. Cells nuclei are in blue. **(B).** Adult HSPCs exhibit a reduced polarization and the AML patient blasts a reverted polarization upon interaction with AML MSCs in the microwells. The quantification of the polarization index indicates a more randomized distribution of the centrosomes of adult HSPCs in interaction with both AML MSCs derived from two AML patients 4 and 5. AML patient blasts demonstrate a reverted polarization with significant number of cells displaying distal positions of their centrosomes during contacts with AML MSCs. Each spot shape denotes a biological replicate (three replicates; n _adult HSPC / HD MSC_ = 88, n _adult HSPC / Patient 4 MSC_ = 84; n _adult HSPC / Patient 5 MSC_ = 135, n _Patient 1 blasts / Patient 4 MSC_ = 73). The mean of the medians is represented by a black bar. Differences between populations were evaluated using a Kruskal-Wallis ANOVA test with P-values of 0.0060, 0.0222 and <0.0001 (**, *, ****).

We took advantage of our recent design of a microfluidic chip that reconstitutes a simplified but transparent bone-marrow in a microfluidic circuit, ie a “bone-marow on-a-chip” (BMoC), in order to investigate cell migration properties^31^. Our BMoC incorporates distinct compartments for stromal cells, supporting long-term culture of osteoblasts, endothelial cells, as well as primary stromal cells from healthy donors or AML patients, all encapsulated in a collagen-fibrin-based hydrogel (*Figure 4.A*). The three-dimensional environment provided by this hydrogel allows hematopoietic cells to visit all compartment and possibly establish contact, detach, or migrate in contact with specific stromal cells. The BMoC design also enables continuous monitoring of live cells, as well as fixation, immunostaining, and high-resolution imaging of fixed samples (*Figure 4.A*).

**Figure 4:**
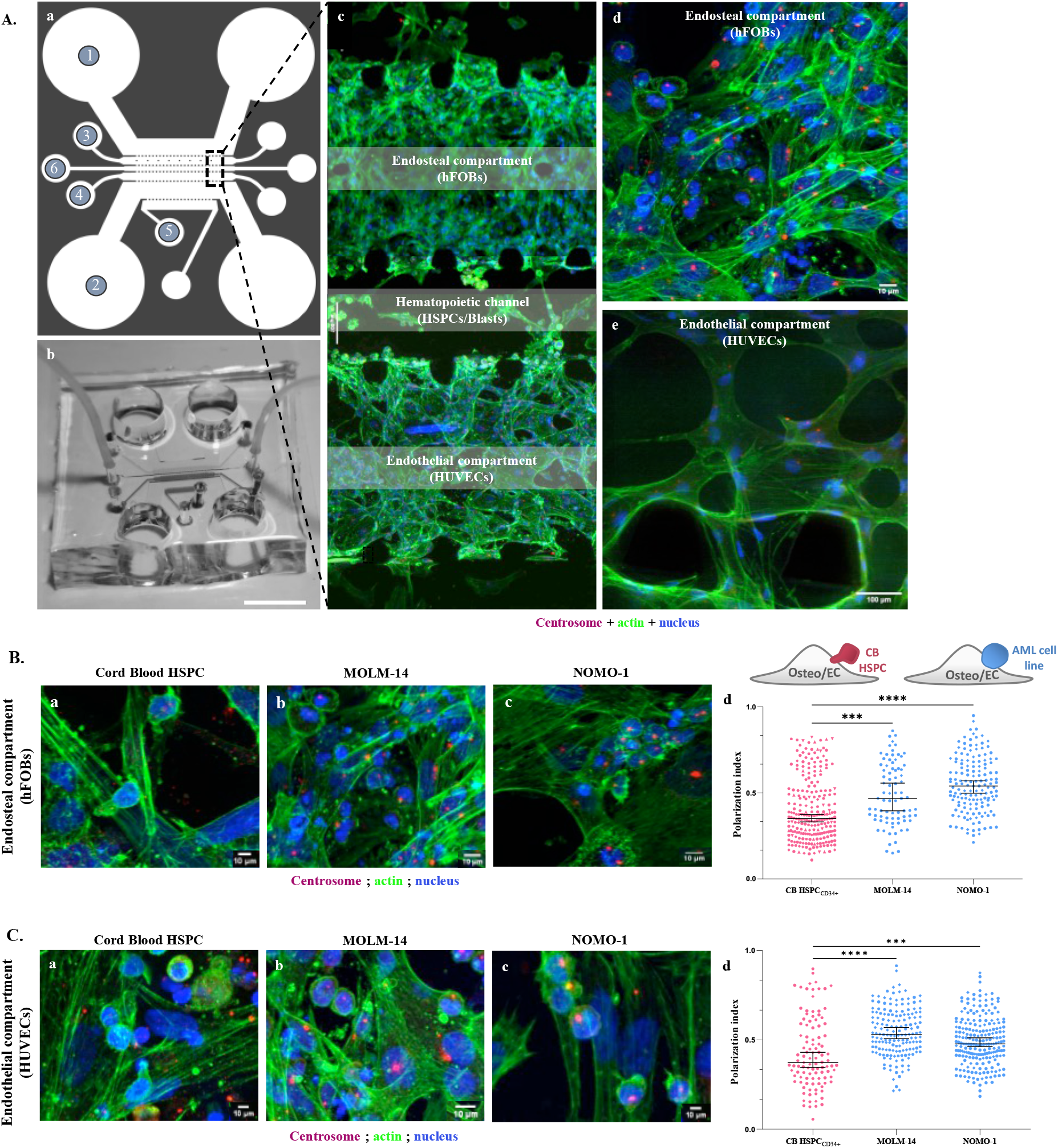
The compartmentalization of the Bone Marrow on Chip (BMoC) promotes both interaction and migration of hematopoietic cells. **(A). (a)**. The microfluidic circuit comprises 6 parallel channels. Channels *(1)* and *(2)* are committed to medium supplementation. Channels *(3), (4)* and *(5)* are dedicated to endosteal (hFOBs), endothelial (HUVECs) and fibroblastic cells (NHLFs), respectively. Endosteal compartment harbours central pillars to decrease traction forces displayed by osteoblasts within the compartment. Subsequently, the work involved using mesenchymal stem cells (MSCs) sorted from either healthy donor or AML patient bone marrow. All stromal cells are loaded encapsulated in a hydrogel composed of collagen and fibrin. The central channel *(6)* is devoted to hosting hematopoietic cells such as hematopoietic stem and progenitor cells (HSPCs), AML cell lines (MOLM-14 or NOMO-1), or AML patient blasts. **(b)**. Photography of the BMoC perfused with colorants to illustrate the position of the distinct compartments. **(c)**. Overall view of the organisation of both stromal compartments visualised via actin labelling (green). **(d)**. Organisation within the endosteal compartment comprising osteoblasts (hFOB) and an AML cell line (NOMO-1). **(e)**. Organization within the endothelial compartment comprising endothelial cells (HUVECs) that aggregate to form a tri-dimensional network reminiscent of tube-like structures. Cell nuclei and centrosomes are displayed in blue and red respectively. **(B).** AML cell lines MOLM-14 and NOMO-1 do not exhibit polarisation upon interaction with osteoblasts in the BMoC. **(a-c)**. CB HSPCs, MOLM-14 and NOMO-1 interacting with osteoblasts (hFOBs) in the BMoC endosteal compartment. Centrosome, actin and nuclei are visualized in red, green and blue respectively. **(d)**. Quantification of the polarisation index of AML cell line 1 (MOLM-14) and AML cell line 2 (NOMO-1) in comparison to CB HSPCs in contact with osteoblasts. Each spot shape denotes a biological replicate (five or three replicates; n _CB HSPCs_ = 240, n _MOLM-14_ = 81; n _NOMO-1_ = 153). The mean of the medians is represented by a black bar. Differences between populations were evaluated using Kruskal-Wallis ANOVA test with P-values of 0.0002 and <0.0001 (***,****). **(C).** AML cell lines MOLM-14 and NOMO-1 do not exhibit polarisation upon interaction with endothelial cells in the BMoC. **(a-c)**. CB HSPCs, MOLM-14 and NOMO-1 interacting with endothelial cells (HUVECs) in the BMoC endothelial compartment. Centrosome, actin and nuclei are visualized in red, green and blue respectively. **(d)**. Quantification of the polarisation index of AML cell line1 (MOLM-14) and AML cell line2 (NOMO-1) in comparison to CB HSPCs in contact with endothelial cells. Each spot shape denotes a biological replicate (three replicates; n _CB HSPCs_ = 106, n _MOLM-14_ = 167; n _NOMO-1_ = 215). The mean of the medians is represented by a black bar. Differences between populations were evaluated using Kruskal-Wallis ANOVA test with P-values <0.0001 and 0.0008 (****,***).

We first tested the ability of HSPCs to polarize in contact with either endosteal (hFOBs) (*Figure 4.B.a-c*) or endothelial (HUVECs) cells (*Figure 4.C.a-c*) in the BMoC. In accordance with our previous observations of confined hematopoietic-stromal cell doublets in microwells, HSPCs appeared capable of polarizing towards both cell types, while AML cell lines exhibited a random polarity (*Figure 4.B.d, 4.C.d*). We then investigated the migration of AML cell lines and patient blasts in compartments containing stromal cells from healthy donors. Remarkably, the cell line exhibited minimal movement, whereas patient blasts demonstrated swift and processive migration tracks (see supplementary movie S1). This observation suggested that long-term culture of cell lines in flasks without physiological anchorage to a physiological niche may have led to a progressive loss of migration properties. This prompted us to shift our focus to primary hematopoietic cells.

We analyzed the migration trajectories of healthy donor derived HSPC (HD-HSPCs) and AML-derived blasts in contact with stromal cells from healthy or AML patients (Figure 5.A). Interestingly, HD-HSPCs exhibited faster and more persistent movements when in contact with AML-derived stromal cells compared to stromal cells derived from healthy donors (*Figure 5.B*). These findings align with our observation in microwells, indicating defective polarization and thus increased ability of healthy HSPCs to detach and move away when in contact with stromal cells derived from AML patients. Furthermore, upon reconstituting the conditions mimicking AML patients (blasts and MSC derived from AML patients), we found that blasts, displayed even more accelerated and extensive movement on AML-derived stromal cells than on healthy stromal cells (*Figure 5.C*). Interestingly, these data showed that the leukemic niche actively promotes the migration of hematopoietic cells, whether they are transformed or not. But they also demonstrated that the combination of leukemic blast and leukemic niche leads to higher motility than any other combination of hematopoietic and stromal cells.

**Figure 5:**
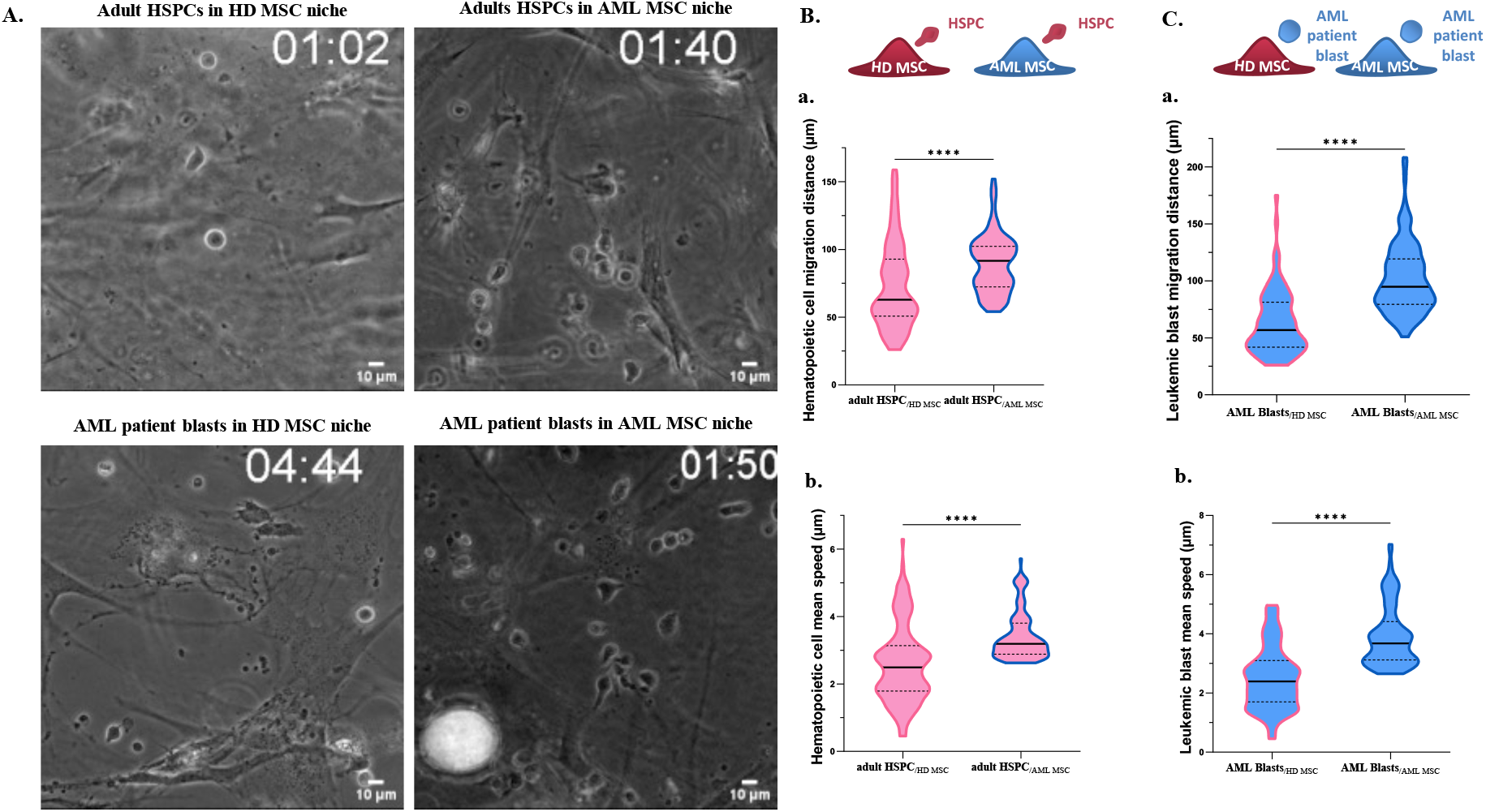
Live imaging within the BMoC enables to decipher increased motility profiles of both adult HSPCs and AML patient blasts in a humanized leukemic stromal compartment. **(A).** Live imaging of healthy HSPC or AML patient blasts within healthy or leukemic compartments. **(a)** Healthy adult HSPCs in an HD MSC compartment. **(b)**. Healthy adult HSPCs in an AML MSCs compartment (derived from AML patient 6). **(c)**. AML patient blasts (AML patient 7) in an HD MSC compartment. **(d)**. AML patient blasts (AML patient 6) in an AML MSC compartment (AML patient 5). **(B).** Quantification of healthy hematopoietic cells (HSPCs) or **(C)**. leukemic blast migration **(a)** and mean speed **(b)** regarding the nature of the compartment (healthy or leukemic MSCs compartments). Both total distances travelled and mean speed, are measured via the TrackMate platform (Fiji) regarding track durations ranging from 20 to 40 minutes. **(B.a-b)**. Tracks of adult HSPCs were delineated from numerous spots within the stromal compartment (n _adult HSPC/ HD MSC_ = 88, n _adult HSPC/ AML MSC_ = 95). **(C.a-b)**. Tracks of leukemic blasts were delineated from numerous spots within the stromal compartment (n _AML blast / HD MSC_ = 140, n _AML blast / AML MSC_ = 116). The mean of the medians is represented by a black bar. Differences between populations were evaluated using a Mann-Whitney test with P-values <0.0001 (****).

## Discussion

In this study, we identified the impaired ability of blasts to engage in interaction and polarization when in contact with the stromal cells of their niche. Initially, we reported the polarization of healthy cord blood-derived HSPCs in contact with osteoblastic or endothelial cell lines^25^. Here, we confirmed that adult HSPCs have retained this property. Interestingly, we found that adult HSPCs could not polarize in contact with leukemic stromal cells. This suggested that the leukemic niche can have a detrimental impact on the structural organization of healthy HSPCs. Conversely, we found that leukemic blasts could not polarize toward either healthy or leukemic stromal cells. So, collectively, our findings demonstrated that the malignancy of either stromal or hematopoietic cells from AML patients hinders polarization of hematopoietic cells. Moreover, they indicated that the combination of a leukemic blast in a leukemic niche fully reverses blast polarity and promotes its migration. This showed that both intrinsic and extrinsic factors are involved in the regulation of the polarization of hematopoietic progenitors, as it has been demonstrated in other tissues^34^. Furthermore, our data showed that both hematopoietic cells and stromal cells from the niche contribute to the morphological transformation of leukemic cells, contributing to the substantial body of evidences indicating the critical role of this interplay in leukemia progression^8^.

Further investigation is warranted to elucidate the molecular pathways that are affected by the leukemic context and lead to the loss of cell polarity. Our work showed the implication of the SDF1-CXCR4 pathway in the polarization and orientation of the division of cord blood-derived HSPCs^25,35^. This pathway being involved in leukemic transformation^36^, it is thus likely to be involved in the reduced contact and defective polarization of leukemic blasts that we observed, as well as the numerous other adhesion pathways engaged in stem cell niche retention^37^.

The causal relationship between polarity defects and AML remains indeterminate, necessitating an exploration of the interrelation between anchorage/polarity and proliferation/differentiation processes. Maintaining the balance between hematopoietic cell expansion and differentiation, a balance that is disturbed in AMLs, likely intricately hinges on cell polarization in interphase and asymmetric cell divisions, as it does in other stem cells^38,39,40^. Hence, further examination is required, along with exploration of polarity together with the identity of progeny cells, through the defective segregation of signals during leukemic cell division^27,29,41^. In addition, cell polarization might be directly coupled to cell quiescence. Indeed, in multiple systems, the AMPK-LKB1 pathway connects cell polarity to the control of cell quiescence, by associating metabolic needs to the organization of cytoskeleton networks^42,43^. It might thus directly link HSC polarization to the balance between quiescence and proliferation^44,28^.

Noteworthy, the devised complementary engineered artificial niches present a methodological approach to quantify the functional behavior of blasts derived from patients, assessing the polarity across different stages of AML, with potential implications for future prognostic evaluations, as well as identification of the exact contribution of the loss of blast polarity to the progression of leukemia. They also constitute a convenient platform to parallelize genomic or chemical screens combined with automated image analysis^45^ in order to reveal the molecular mechanism underlying the defective polarization of leukemic cells and identify chemical strategies to revert it.

## Methods

### 1. Hematopoietic Cell Purification

#### 1.1 Collection and Processing of Cord Blood and Bone Fragments

Human umbilical cord blood cells were obtained from a compliant institution with ethical review board approval. Cord blood bags, provided by the Cord Blood Bank of Public Assistance – Hospital of Paris (AP-HP), were used in coordination with the Biological Resources Centre authorized by the French Cord Blood Network. Leftover hip bone fragments from adult donors to obtain adult HSPCs were collected following hip replacement surgery at Lariboisière Fernand-Widal AP-HP. Mononuclear cells were isolated using a density gradient, separating them from plasma, granulocytes, and erythrocytes using a lymphocyte separation medium (euro Bio, AbyCys). CD34+ cells from cord blood or adult donors were sorted using microbeads conjugated to monoclonal mouse anti-human CD34 antibodies (Miltenyi Biotec MACS) through positive magnetic bead selection on LS columns.

#### 1.2 AML Patient Blast Samples

Blasts from AML patients were obtained from consenting individuals registered in an ongoing clinical registry at Saint Louis Hospital (THEMA, IRB approval no. IDRCB 2021-A00940-41). Cytogenetic analyses, karyotyping, and fluorescence in situ hybridization studies were conducted, as part of Saint Louis hospital standard of care clinical practice, along with genetic profiling using a Next Generation Sequencing (NGS) panel targeting 94 myeloid genes (*ACD, ALDH2, ARID2, ASXL1, ASXL2, BCOR, BCORL1, BRAF, BRCA1, BRCA2, BRCC3, CALR, CBL, CCND1, CCND2, CDKN2A, CDKN2B, CEBPA, CHEK2, CREBBP, CSF3R, CSNK1A1, CTCF, CUX1, DDX41, DNM2, DNMT3A, EIF6, EP300, ERCC6L2, ETNK1, ETV6, EZH2, FLT3, GATA2, HRAS, IDH1, IDH2, IKZF1, IKZF5, IRF1, JAK2, JAK3, KDM5A, KDM6A, KIT, KMT2A/MLL, KMT2D, KRAS, LUC7L2, MBD4, MECOM, MGA, MPL, MPO, MYC, NF1, NPM1, NRAS, PDS5B, PHF6, PPM1D, PRPF8, PTEN, PTPN11, RAD21, RIT1, RUNX1, SAMD9, SAMD9L, SBDS, SETBP1, SETD1B, SF1, SF3B1, SH2B3, SMC1A, SMC3, SP1, SRP72, SRSF2, STAG2, TERC, TERT, TET2, TP53, TRIB1, U2AF1, U2AF2, UBA1, UBE2A, WT1, ZNF687, ZRSR2*) to determine mutation status. Mononuclear cells were extracted from bone marrow samples using Ficoll-Paque™ PLUS (GE Healthcare #17-1440-02), followed by red blood cell lysis using a red blood lysis buffer (Sigma #R7757).

#### 1.3 Hematopoietic Cell Culture Conditions

The culture of hematopoietic cells, including cord blood/adult HSPC and AML patient blasts, involved maintenance in RPMI supplemented with fetal bovine serum (FBS), 1% of antibiotics/antimycotics (100 U/ml Penicillin, 100 µg/ml Streptomycin, 0,25 µg/ml Fungizone, (Gibco #15240062), and specific cytokines.

### 2. Healthy donor and AML patient MSCs

Healthy donor MSCs were extracted from hip bone fragments of adult donors, while AML MSCs were cultured from bone marrow samples of AML patients. Before extracting mononuclear cells, a substrate of the bone marrow sample was cultured enabling MSCs growth in α-MEM ((StableCell Minimum Essential Medium Eagle – α modification, Sigma #M6199) supplemented with human Platelet Lysate (hPL, StemCell#06960) and 1% antibiotics/antimycotics.

### 3 Cell lines culture

HUVECs (Lonza #00191027) and hFOB 1.19 (ATCC CRL-11372TM) were cultured in EBM-2 (Lonza #CC-3162) supplemented with EGM™ SingleQuots (Lonza #CC-4176), and in Ham’s F12 DMEM (Dulbecco’s Modified Eagle’s Medium, 2,5 mM L-Glutamine/GlutaMAX™) supplemented with 10% FBS and 1% antibiotics/antimycotics, respectively. MOLM-14 and NOMO-1 AML cell lines were cultured in RPMI (Roswell Park Memorial Institute 1640 medium) supplemented with 10% of FBS and 1% of antibiotics/antimycotics.

Mycoplasma testing was conducted regularly, and cultures were maintained in a controlled environment.

### 4. Microwells microfabrication and assembly

#### 4.1 Polyacrylamide microwells microfabrication

Microwell specifications, including shape, size, and layout, are designed using CleWin software to be subsequently transferred to a quartz photomask coated with a chromium layer (Toppan Photomask). They are fabricated on a silicon wafer. A negative mold of the silicon wafer is made in PDMS (polydimethylsiloxane SYLGARD184™ - Dow Corning Kit). From a second positive PDMS stamp of microwells, stamps of 0.5×0.5cm are cut.

Glass coverslips of 20×20 µm are coated with silane (2% (3-Trimethoxysil) propyl-methacrylate (Sigma #M6514)), and 1% acetic acid (Sigma-Aldrich #818755) in ethanol.

After a plasma treatment, the PDMS stamp is placed on a silanized coverslip pretreated with 2% (3-Trimethoxysil) propyl-methacrylate (Sigma #M6514)), and 1% acetic acid (Sigma-Aldrich #818755) in ethanol. A mixture containing 20% of a solution of polyacrylamide (acrylamide/bis-acrylamide 40% 37.5 (Euromedex #EU0062-B)), 1% of a photo-initiator (2-hydroxy)2-methylpropiophenone (MPP - Sigma-Aldrich #405655), 0.5% ammonium persulfate (APS, Thermo Scientific #17874), 0,5% tetra-methyl-ethylene-diamine (TEMED, Sigma #T9281) in MilliQ water is introduced between the PDMS and the coverslip by capillary forces. The mounted system is irradiated at 23mJ/cm^2^ for 5 minutes to cure the polyacrylamide gel and allow the stamp removal. For sterilization, microwells are irradiated in 70% ethanol, then rinsed in PBS overnight to remove any residual contaminants.

#### 4.2 Microwells assembly

Microwells were coated with fibronectin (10µg/ml, Sigma #F1141). After PBS rinse, stromal cells [hFOB - HUVEC - HD MSC - AML patient MSC] were added in their adequate media. After incubation, hematopoietic cells [CB/adult CD34+ HSPC – MOLM-14 – NOMO-1 – AML patient blasts] were loaded and incubated in their regular media.

### 5. Bone Marrow on Chip Assembly

#### 5.1 Microfabrication and Outsourcing

The chip design as reported in previous lab works ^31 25^, inspired by the Noo Li Jeon laboratory, faced limitations in homemade manufacturing, leading to outsourcing to Biomimetic Environment On Chip company. The chips underwent UV treatment for sterilization before use.

#### 5.2 Hydrogel Composition and Cell Loading

A solution of rat tail collagen type I (Ibidi #50201) at 1,6 mg/ml, fibrinogen at 2 mg/ml (Sigma #F3879), and thrombin at 1U/ml (Sigma # T6884) compose the hydrogel encapsulating stromal cells. 2.10^5^ HUVECs or hFOBs or HD MSCs or AML MSCs are loaded in the gel within the dedicated channels.

The BMoC is then incubated for gelation at 37°C for approximately 45 minutes. Regular mediums of the stromal cells are injected into the medium inlets.

When the stromal cell compartments are functional (∼48h) cord blood and adult CD34^+^ HSPCs, AML cell lines, or AML patient blasts can be loaded in the central channel of the chip. All medium inlets are subsequently filled with the regular medium of the hematopoietic cells.

### 6. Immunostaining

Cells within microwells or chips were fixed (4% paraformaldehyde (16% PFA, Delta Microscopies #D15710)), permeabilized (0.1% Triton X-100 (Sigma-Aldrich)), and blocked (3% BSA (Bovine Serum Albumin, Sigma-Aldrich #A7030) and 0.1% Tween (Sigma-ldrich)). Primary antibodies included polyclonal rabbit anti-pericentrin (Abcam #ab4448) at 1/600, and monoclonal mouse anti-pericentrin (Abcam #ab47654) at 1/200. Cross-absorbed secondary antibodies Alexa Fluor™ 568nm- and 647nm-conjugated goat anti-rabbit and anti-mouse (Life Technologies #A11004 and #A21240) were used at 1/600. F-actin was labelled with Alexa Fluor 488-conjugated phalloidin (Sigma-Aldrich #A12379). Nucleus localization is stained using DAPI (Sigma-Aldrich #D9542). The coverslips containing microwells are mounted in mowiol after the removal of polyacrylamide microwell walls (Sigma-Aldrich #81381). The BMoCs are mounted in mowiol added within their inlets and outlets.

### 7. Microscopy

#### 7.1 Live and immunofluorescence imaging

Time-lapse imaging for BMoC required the use of the Nikon Biostation IM-Q Cell-S2 microscope (Institute of Research of Saint Louis platform). After loading with hematopoietic cells, the devices are placed in an enclosed chamber at 37°C with controlled humidity and normoxic conditions to be imaged every 2 minutes. Image processing is performed with 20X 0.80-NA air objective and binnings of 2×2, using the Biostation interface 2.1. Images of fixed samples are captured using a Nikon Ti-eclipse microscope equipped with a spinning disc (Yokogawa-CSU-X1) and a 100X 1.40-NA oil objective, together with an electron-multiplying charge-coupled device camera (CCD camera Photometrics-Evolve512). The acquisition software used is MetaMorph. Z-stacks of 0.5μm thickness were acquired using a 2×2 binning.

#### 7.2 Polarization index measurement

Centrosome position was measured using a MACRO via the Fiji interface. First, the centrosome position is manually marked. Second, the membrane zone of the cell of interest where it interacts with the stromal cell is marked to record the distance to the centrosome (distance d). Finally, a third point representing the localization of the tip of the cell is selected as the point in the cell furthest away from the previous one. The polarisation index is calculated as the ratio of the distance from the centrosome to the contact zone, to the total length of the cell (d/D).

#### 7.3 Cell tracking

Cell tracking is performed using TrackMate7 an open-source platform, distributed in Fiji, for single-particle tracking, data visualization and editing results.^46^ Relying on three filters on tracks, track duration, total displacement and mean speed are set to compare tracks displaying the same time.

### 8. Statistics

All of the statistical analyses were carried out using GraphPad Prism 9 software. A Shapiro-Wilk test is used to assess the normality of the data distribution and to guide the selection of appropriate statistical tests. Differences between populations are assessed using non-parametric methods such as Kruskal-Wallis ANOVA and Mann-Whitney t-test for non-normally distributed samples, or parametric tests such as unpaired t-test and one-way ANOVA for normally distributed samples. All data are presented as superplot groups with different biological replicates, each represented by a distinct spot shape.

## Acknowledgements

We express our gratitude to A. Puissant and R. Itzykson (Saint Louis Hospital, U944, Université Paris Cité) for their valuable scientific insights. Our appreciation extends to Niclas Setterblad from the Saint-Louis Research Institute Core Facility at Saint-Louis Hospital.

This work was supported by the European Research Council (Consolidator Grant 771599 to M.T., the INCa (Institut National du Cancer PRTK 2022-192 to L.B.) and the ATIP-Avenir 2022 program (Inserm and Ligue Nationale Contre le Cancer N° 23AA019-00 to L.B.), by the Bettencourt-Schueller Foundation, the Emergence program of the Ville de Paris and the Schlumberger Foundation for Education and Research. The Integrated Cancer Research Center, “SiRIC InsiTu: Insights into Cancer: From Inflammation to Tumor,” funded this study, under grant number INCa-DGOS-INSERM-ITMO Cancer_18008.

This research also received financial support from a doctoral school contract (to K.S.) provided by the University of Paris Cité, as well as from The National Center for Precision Medicine in Leukemia (THEMA) of Saint Louis Hospital (Paris, France) and Bristol Myers Squibb (BMS).

## Authorship Contributions

**K.S**.: conceptualization, formal analysis, validation, investigation, visualization, methodology, writing-original draft. **B.V**.: conceptualization, methodology, supervision, investigation. **L.Bo**.: conceptualization, methodology, investigation. **L.K**.: resources. **H.P**.: resources. **C.C**.: resources. **R.M**.: resources. **S.F**.: resources. **P.C**.: resources. **E.K**.: resources. **A.J**.: resources. **E.R**.: resources. **R.N**.: resources. **L.Bl**.: supervision, funding acquisition. **L.Be**.: resources, revision of the original draft. **M.T**.: conceptualization, supervision, funding acquisition, investigation, methodology, writing-original draft, project administration.

## Supplementary Figure: AML patient cytogenetic and molecular characteristics

**Table 1:** AML patient clinical characteristics. Anonymized clinical data of AML patient ages, genders, disease statuses, time sampling, and nature of the extracted cells.

| Patient | Age | Gender | Disease Status | Time sampling | Extracted cell type |
| --- | --- | --- | --- | --- | --- |
| Patient n°1 | 49 | Male | De novo AML | Primary Refractory | *Blasts* |
| Patient n°2 | 57 | Female | De novo AML | Complete Remission | *Blasts* |
| Patient n°3 | 44 | Male | De novo AML | Complete Remission | *Blasts* |
| Patient n°4 | 27 | Male | De novo AML | Complete Remission | *MSCs* |
| Patient n°5 | 20 | Male | De novo AML | Complete Remission | *MSCs & Blasts* |
| Patient n°6 | 21 | Male | De novo AML | Complete Remission | *MSCs* |
| Patient n°7 | 59 | Female | De novo AML | Not available | *Blasts* |

**Table 2:** AML patient cytogenetic profiles. Anonymized clinical data of AML patient donors of either AML MSCs or blasts. Bone marrow blast percentage, karyotype, mutated genes and AML classification (ELN classification 2022).

| Patient | Bone Marrow blasts % | Karyotype | Mutated genes | ELN Classification 2022 |
| --- | --- | --- | --- | --- |
| Patient n°1 | 67 | normal | *IDH2; DNMT3A; BCOR* | Intermediate |
| Patient n°2 | 98 | normal | *NPM1; DNMT3A; WT1* | Favorable |
| Patient n°3 | 92 | normal | *FLT3-ITD; CEBPa* | Favorable |
| Patient n°4 | 52 | 45,X,-Y,t(8;21)(q22;q22)[22] | *FLT3-ITD; MGA* | Favorable |
| Patient n°5 | 60 | 47,XY, inv(16)(p13q24),+21[20] | No additional mutation | Favorable |
| Patient n°6 | 82 | normal | *NRAS; PTPN11; CEBPa; GATA2* | Intermediate |
| Patient n°7 | 35 | 46,XX,t(11;19)(q23;p13)[20] | *NRAS; KRAS* | Adverse |

